# Genome-wide measurement of local nucleosome array regularity and spacing by nanopore sequencing

**DOI:** 10.1101/272526

**Authors:** Sandro Baldi, Stefan Krebs, Helmut Blum, Peter B. Becker

## Abstract

The nature of chromatin as regular succession of nucleosomes has gained iconic status. However, since most nucleosomes in metazoans are poorly positioned it is unknown to which extent the bulk genomic nucleosome repeat length (NRL) reflects the regularity and spacing of nucleosome arrays at individual loci. We describe a new approach to map nucleosome array regularity and spacing through sequencing oligonucleosome-derived DNA by Illumina sequencing as well as emergent nanopore-technology. This revealed modulation of array regularity and NRL depending on functional chromatin states independently of nucleosome phasing and even in unmappable regions. We also found that nucleosome arrays downstream of silent promoters are considerably more regular than those downstream of highly expressed ones, despite more extensive nucleosome phasing of the latter. Our approach is generally applicable and provides an important parameter of chromatin organisation that so far had been missing.

## Introduction

Eukaryotic DNA is stored as chromatin, which consists of repeating nucleosome units, each containing 147 bp of DNA wrapped around histone octamers separated by linker DNA ^1^. Chromatin is not only important for storing the genetic information, but also critically involved in regulating the accessibility of the DNA to transcription factors, RNA and DNA polymerases, and other proteins involved in gene regulation, DNA replication and repair ^2-4^.

The exact position of nucleosomes relative to the underlying DNA sequence – referred to as nucleosome *positioning* – has important consequences for the recognition of DNA sequence motifs by regulatory proteins ^2,3,5^. A nucleosome that organizes an identical stretch of DNA in the majority of nuclei of a population of cells is *well-positioned.* If several nucleosomes in an array are positioned with respect to the underlying DNA sequence, we call them *phased*. Prominent nucleosome phasing occurs adjacent to boundaries, such as nucleosome-free regions of promoters or tightly bound proteins, such as insulators ^2,3^.

Neighboring nucleosome core particles are separated by linker DNA, which provides binding sites for linker histones, remodeling factors and other DNA binding proteins. If linker lengths within a nucleosomal array are relatively constant, then the array is *regular* and the spacing of nucleosomes defines a *nucleosome repeat length (NRL*). Average NRLs in a sample of nuclei can be derived from bulk chromatin by partial cleavage of linker DNA with Micrococcal Nuclease (MNase). The fact that average NRLs varies depending on species, cell type, functional chromatin state and chromosomal position^6-11^ shows that a continuum of chromatin structures is compatible with genome functions. The average NRL of a chromatin sample does not inform about the spacing and regularity at individual loci. Indeed, numerous observations suggest that the regularity of nucleosomal arrays varies considerably along the chromosome. For example, inactive heterochromatic domains that package repetitive DNA show extensive regularity, while the spacing of nucleosomes in euchromatin is more variable ^12-14^. Conceivably, the regular nucleosome spacing is disrupted in active chromatin due to the many remodeling events associated with the usage of chromatin as a template for transcription, replication and repair processes.

One fundamental problem for measuring array regularity and nucleosome spacing at specific sites along the chromosome arises from the fact that most nucleosomes in metazoans are poorly positioned ^6,7,14,15^ as the histone octamers do not organize exactly the same stretch of DNA in every cell, but may be translationally shifted. Regular nucleosome arrays that are translationally shifted in different cells will appear irregular in MNase-seq experiments that map the positions of individual nucleosomes in a population of cells (Fig. 1a, yellow shaded). Thus, in regions with poor nucleosome positioning, we cannot determine array regularity. A method is needed that measures the spacing between neighboring nucleosomes on individual DNA molecules. In principle, this can be achieved by tuning the digestion of chromatin with MNase so that not only mononucleosomes are obtained, but a series of fragments derived from di-, tri-, tetra-, pentanucleosomes and beyond. Running these fragments on a gel reveals a typical ‘MNase ladder’, where the average NRL can be determined from the fragment lengths and the sharpness of the bands serves as a measure for the regularity of the underlying chromatin. Determining the regularity and NRL at an individual locus traditionally requires cumbersome Southern blotting and hybridization of the MNase ladder with a locus-specific probe. Conceivably, by sequencing an MNase ladder, it should be possible to map the fragments to genomic regions and to measure the regularity of the underlying chromatin through the size distribution of the mapped fragments (Fig. 1a, blue shaded). Very regular chromatin should yield fragments with discrete peaks in the size-distribution, representing mono-, di-, tri-, tetranucleosomal etc. fragments. Irregular chromatin, on the other hand, should show a less discrete size distribution. Importantly, such type of measurement would determine the distances between neighboring nucleosomes independently of their phasing. Although earlier studies have reported on the sequencing of MNase ladders ^16,17^, they concentrated mostly on subnucleosomal particles, as conventional high-throughput sequencing techniques do not yield enough reads for long fragments, especially not over a broad range of fragment sizes. Here, we circumvented this problem by using new nanopore-based sequencing technology, which combines relatively high sequencing depth with complete sequencing of long fragments over a wide size range ^18^. As a complementary approach we harnessed established Illumina sequencing for genome-wide NRL determination. This allowed us to determine array regularity and linker length throughout the *Drosophila* genome and even in unmappable regions, and to reveal that regularity is inversely correlated with transcriptional activity. We refer to our approach as Array-seq to indicate that we are sequencing fragments representing nucleosome arrays rather than mononucleosomes.

**Fig. 1.**
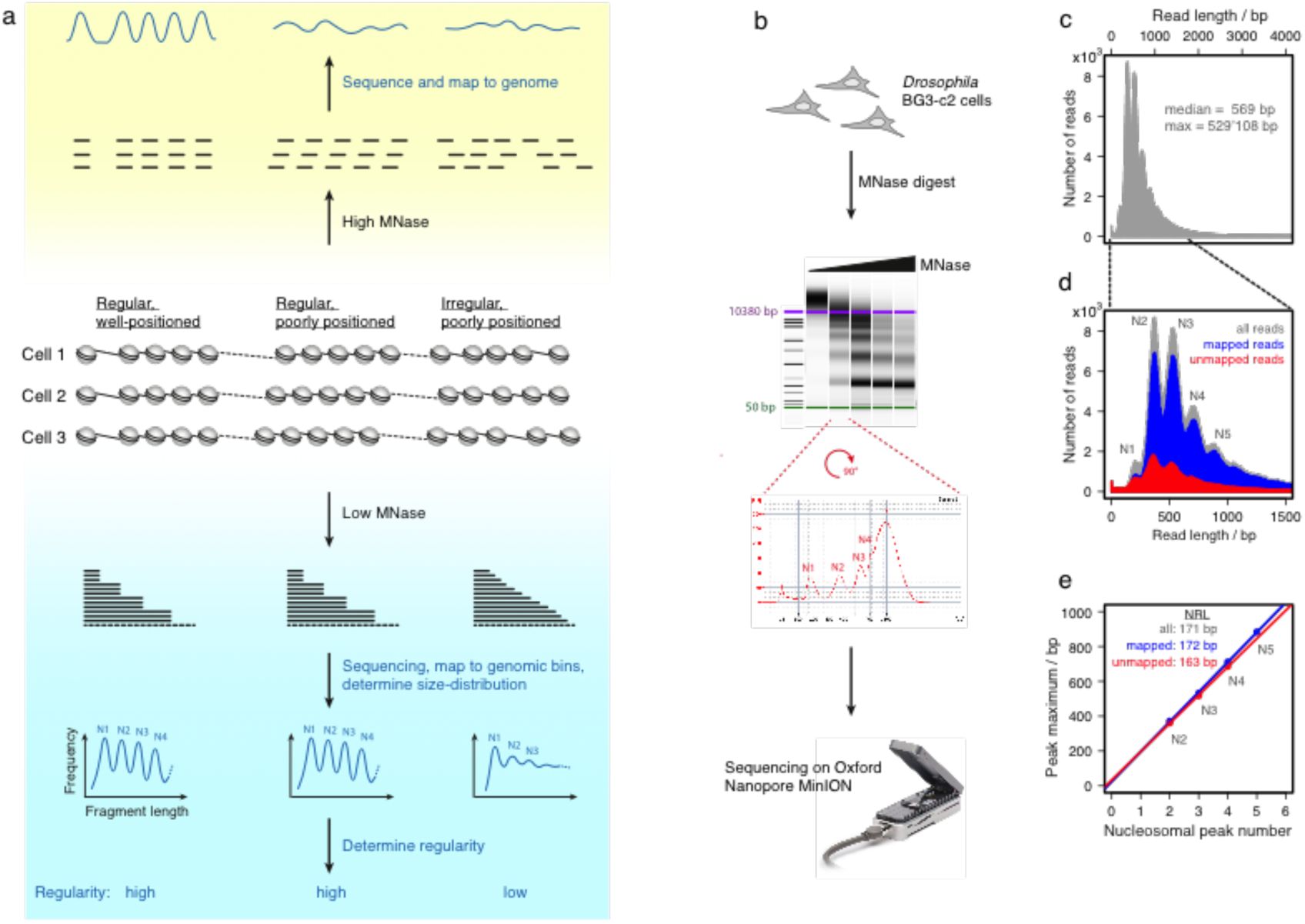
Outline and initial characterization of the Array-seq strategy. A, Schematic representation of MNase-seq strategies. Classic MNase-seq of mononucleosomes (yellow shaded) fails to reveal nucleosome spacing and chromatin regularity in regions with poorly positioned nucleosomes. Limited MNase digestion followed by sequencing (blue shaded) preserves the connectivity between neighboring nucleosomes and therefore allows determining chromatin regularity and linker length even in the absence of well-positioned nucleosomes. **b**, Overview of an Array-seq experiment. The MNase-digested DNA is electrophoresed on a Bioanalyzer device and the sample with the appropriate digestion degree chosen for sequencing on an Oxford Nanopore Minion sequencer. **c**, Size distribution of 796,070 sequenced MNase fragments derived from Drosophila BG3-c2 cells. **d**, Size distribution of fragments below 1500 bp for all sequence reads (grey), the subset of reads that could be mapped to the Drosophila genome (blue), and the unmappable reads (red). N_1_, N_2_ …. indicate the mono-, di-…nucleosomal peaks. **e**, The nucleosome repeat length (NRL) for the sequence reads calculated by determining the slope of the straight line fitted to the di-, tri-, tetra- and pentanucleosomal peak positions of the size distributions.

## Results

### Nanopore sequencing of an MNase ladder

To obtain information about array regularity and linker length in the absence of well-positioned nucleosomes, we digested chromatin from *D. melanogaster* BG3-c2 cells with different amounts of MNase and purified the DNA (Fig. 1b).

From this digestion series we chose an amount of MNase that shows defined laddering and still a large proportion of longer DNA fragments corresponding to oligonucleosomes. DNA derived from this MNase ladder was then sequenced on an Oxford Nanopore MinION device, which yielded a total of 796,070 reads. Although we obtained some very long reads (up to 500 kb), the majority of them had a length below 1000 bp (Fig. 1c). The size distribution of the bulk, unmapped reads suggests defined peaks for di-, tri-, tetra- and pentanucleosomal fragments (Fig. 1d). Notably, mononucleosome-sized fragments are almost completely absent, although they are clearly present in the MNase-digested DNA (Fig. 1b). Presumably, this is because the nanopore device has difficulties to classify short DNA sequences and, by default, discards most of them. However, mononucleosomal fragments are not important for the determination of array regularity and linker length in our approach, as they contain no information about nucleosome spacing. The apparent overrepresentation of long fragments in the Bioanalyzer track of the MNase ladder compared to sequenced reads (Fig. 1b vs. Fig. 1d) is largely explained by the fact that the Bioanalyzer gel electrophoresis device displays DNA mass rather than molecule numbers. The majority (79%) of all sequence reads could be mapped to the *D. melanogaster* genome. The size distribution of the mapped reads was very similar to the one of the total reads. The maxima of the di-, tri-, tetra- and pentanucleosomal peaks were used to determine the average nucleosomal repeat length (NRL) for the sequenced MNase ladder by measuring the slope of the straight line fitted through the peak positions ^19^ (Fig. 1d). For the mapped reads, we obtained an average NRL of 172 bp, which corresponds to a linker length of about 25 bp. This is a bit shorter than the 26-50 bp that was previously reported for NRLs in flies and *Drosophila* cells ^20-22^, but it is known that linker length depends on cell type and genomic position ^6,7,20^.

### Array regularity differs widely for chromatin states

Next, we wanted to know whether array regularity and nucleosome spacing varied depending on the functional state of the genomic region. To do this, we took advantage of the 9-state chromatin model that was previously established for BG3-c2 cells ^23^. This model associates each genomic location with one of nine different states, which are based on combinatorial patterns of histone modifications. These states mostly correspond to distinct classes of transcriptional activity, as they are associated, for example, with active promoters, transcriptional elongation or enhancer signatures, but also with heterochromatin and polycomb-dependent repression. When the Array-seq reads were sorted according to the chromatin state they mapped to, distinct differences in the size distributions emerged (Fig. 2a, second replicate in Supp. Fig. 1a). Some states separate defined oligonucleosomal peaks very clearly, such as state 2 (transcriptional elongation), states 7 and 8 (heterochromatin), and state 9 (extensive silent domains), suggesting a high degree of array regularity with an even spacing between adjacent nucleosomes. States 1 and 3 (active promoters and enhancers, respectively), on the other hand, showed a much more blurred size-distribution, probably due to nucleosome-free regions and bound transcription factors in promoters and enhancers that disrupt the nucleosomal arrays. To quantify the degree of regularity in the size-distribution diagrams, we analyzed them by spectral density estimation (Fig. 2b). The spectral density plots of the different chromatin states contain a peak corresponding to a periodicity of about 180 bp, which represents the contribution of the NRL to the overall signal. The height of this peak is a measure for the regularity of the analyzed chromatin and can be used as a “regularity score” (Fig. 2c). To rule out any effect on the apparent array regularity due to unequal numbers of reads obtained for the different states, we repeated the analysis with equal read numbers for each state (Supp. Fig. 2a&b). However, this did not affect the outcome in any major way. The exact positions of the maxima of the nucleosomal peaks in the size-distribution diagrams were used to calculate the nucleosome repeat length for the different chromatin states (Fig. 2d). The repeat length is relatively short in gene bodies of actively transcribed genes (state 2, 168 bp), and longest in heterochromatin (state 7, 177 bp). This fits well with earlier observations that active transcription seems to decrease linker length and array regularity ^7,9,14,22,24^.

**Fig. 2.**
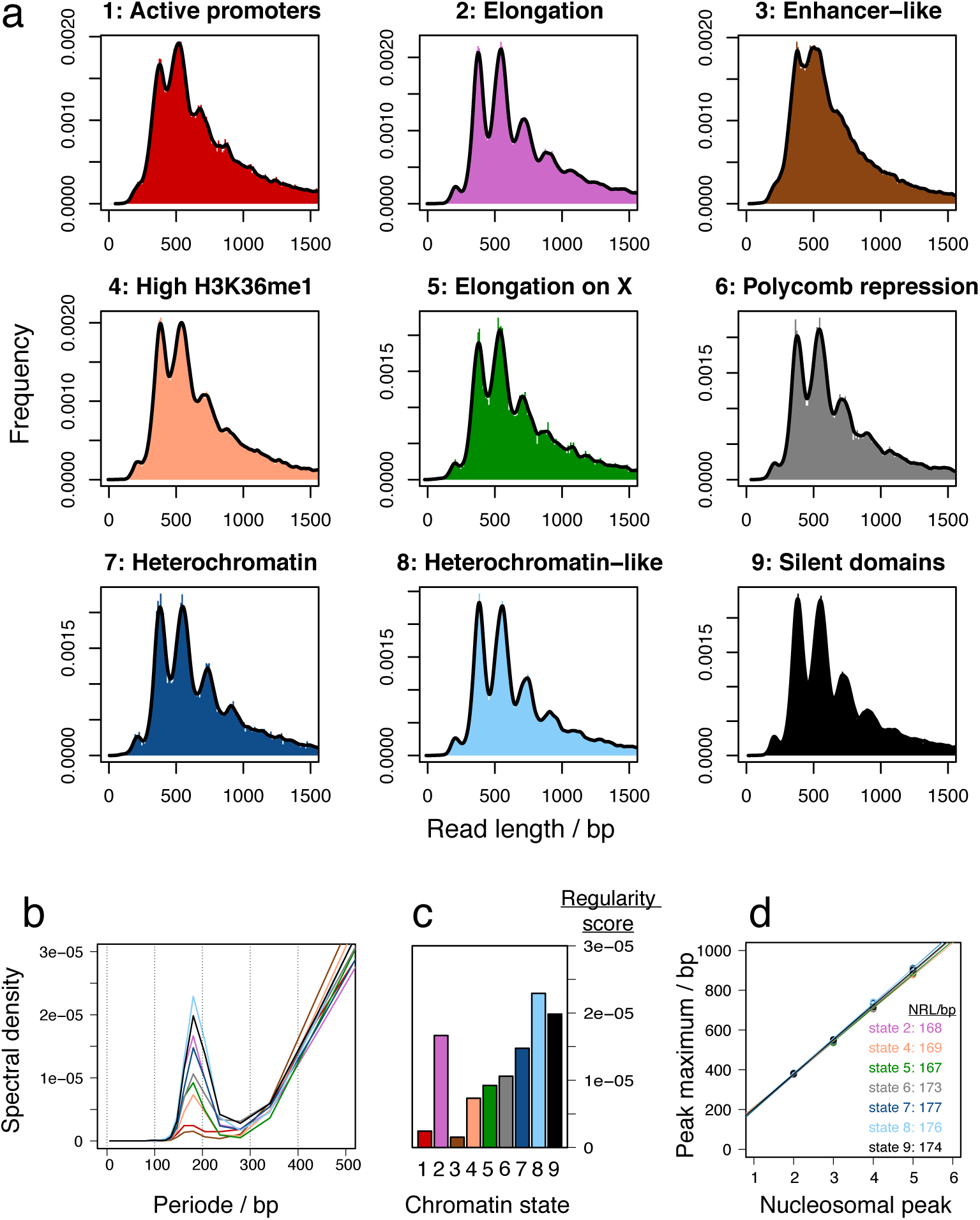
Array-seq analysis of BG3-c2 chromatin. A, Size distributions of Array-seq reads mapping to the different states from the 9-state chromatin model **b**, Spectral density plot of the size distributions in (**a**). The peak at a period of about 180 bp corresponds to the contribution of the nucleosome repeat length to the overall signal. The height of this peak is a measure for the regularity of the underlying chromatin. **c**, Regularity scores for the different chromatin states. Regularity scores correspond to peak heights in the spectral density plot in (**b**). **d**, NRLs for the different chromatin states as determined by the slopes of the fitted lines through the N_2_-N_5_ peaks of the read size distributions.

### Transcription reduces array regularity

Classically, nucleosome spacing has often been analyzed at the 5’ end of expressed genes, where well-positioned nucleosomes can be readily mapped in MNase-seq experiments ^6,7,14,25,26^. With Array-seq it is possible to determine array regularity also in the absence of well-positioned nucleosomes throughout the gene. We determined the size distribution of fragments mapping to gene bodies of either highly or lowly expressed genes (Fig 3a, second replicate Supp. Fig. 1b). First, the NRL is reduced from 173 bp in non-expressed genes to 164 bp in highly expressed ones. Interestingly, the distribution of fragments mapping to highly expressed genes is more blurred than the one mapping to silent genes, presumably due to nucleosome disruption by RNA polymerase ^27^ (Fig. 3a,b). Silent genes show also a slightly increased regularity compared to the genomic average and are comparable in regularity to the heterochromatic and silent chromatin domain states 8 and 9.

**Fig. 3.**
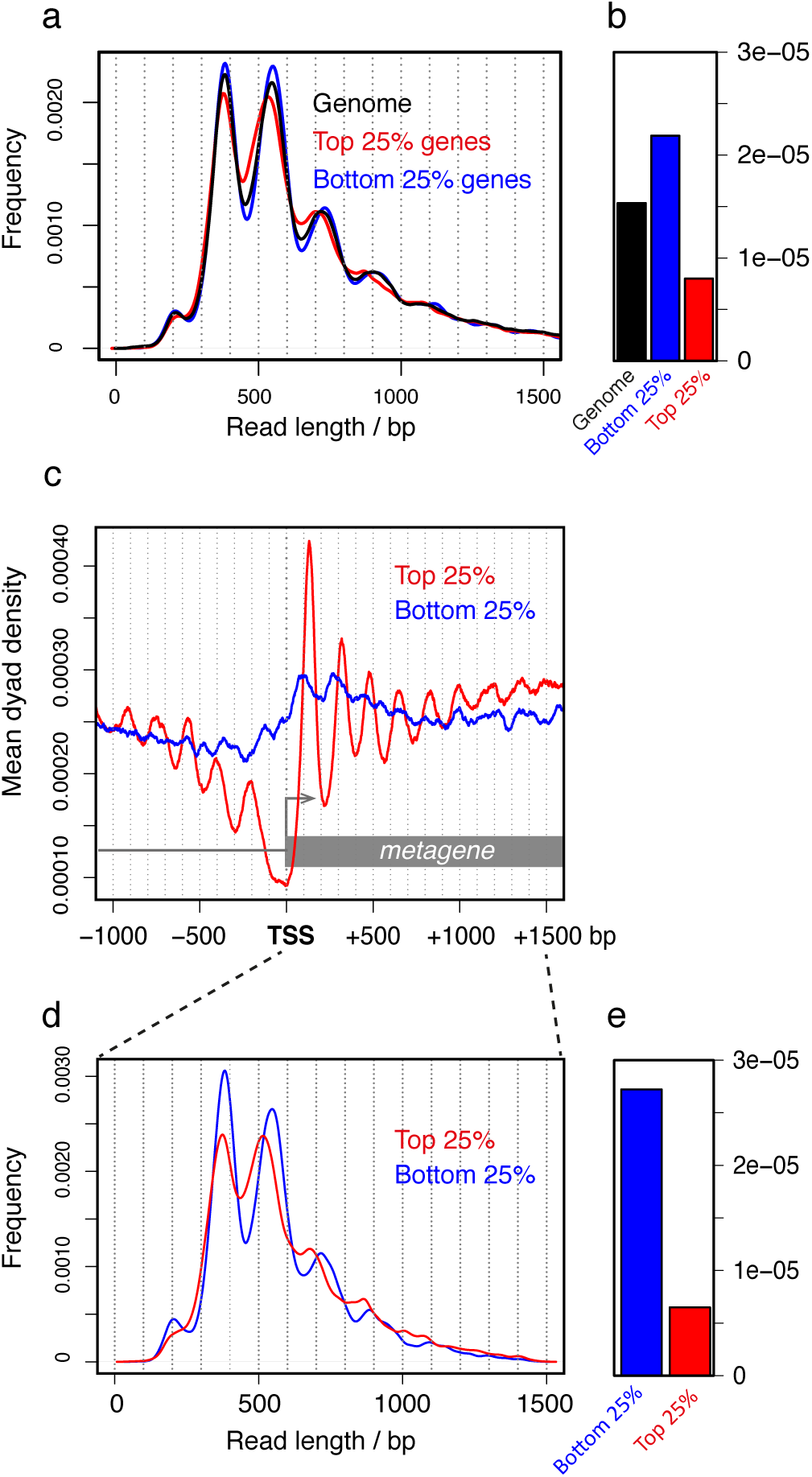
Nucleosome array regularity as a function of transcription activity. A, Size distributions of Array-seq reads from BG3-c2 cells mapped to the whole genome (black), the gene bodies of the 25% least expressed genes (bottom 25%, blue), or the gene bodies of the 25% most expressed genes (top 25%, red). **b**, Regularity scores for the genes grouped in (**a**). **c**, Metagene analysis on mononucleosome dyad densities from BG3-c2 cells. Densities are aligned to the transcription start sites (TSS) of the bottom 25% (blue) or top 25% expressed genes (red). **d**, Size distributions of Array-seq reads mapping to first 1500 bp downstream of TSS of either the bottom 25% (blue) or top 25% expressed genes (red). **e**, Regularity scores for the samples grouped in (**d**).

Superficially, these findings appear counterintuitive, as the 5’ end of active genes are well known for their regularly arrayed nucleosomes ^6,7,14,15,25,26^. Indeed, when we mapped mononucleosomal fragments from BG3-c2 cells and aligned the dyad densities at transcriptional start sites (TSS), well-positioned and evenly phased nucleosomes were observed at highly expressed genes, but not at silent ones (Fig. 3c). Conceivably, chromatin may be more regular in highly expressed genes just downstream of the promoter. However, this is not the case, as when we analyzed the size distribution of Array-seq reads that map to the first 1.5 kb downstream of TSS, we still score a higher regularity in the silent genes compared to the highly expressed ones (Fig. 3d,e) – in fact, the difference is more pronounced than for the whole gene body. This demonstrates that silent genes have actually a high degree of regularity, a conclusion that could not be reached by classic mononucleosome mapping because of the translational shifts of the arrays in inactive genes in the absence of an instructive boundary ^28^.

### Array-seq reveals array regularity and linker length even in unmappable regions

A major obstacle for the analysis of classical MNase-seq experiments is that large parts of metazoan genomes are comprised of repetitive sequences that are only poorly annotated and mostly not included in genome assemblies, precluding the mapping of sequenced reads to these regions. Array-seq, on the other hand, does not strictly depend on the mappability of the reads, since sequences can be categorized as, for example, containing heterochromatic DNA repeats, even if their precise location is unknown (Fig. 4a). We sorted Array-seq reads that contain high proportions of the heterochromatic AATAACATAG satellite repeat ^29,30^ and compared their size distribution to the one of the same number of reads derived from the mappable genome (Fig. 4b,c). Interestingly, the heterochromatin-derived reads showed a very high degree of regularity, separating even the hepta-nucleosomal peak. Accordingly, the regularity score of the AATAACATAG-repeat-containing heterochromatin is much higher than the one derived from the mappable control reads (Fig. 4d).

**Fig. 4.**
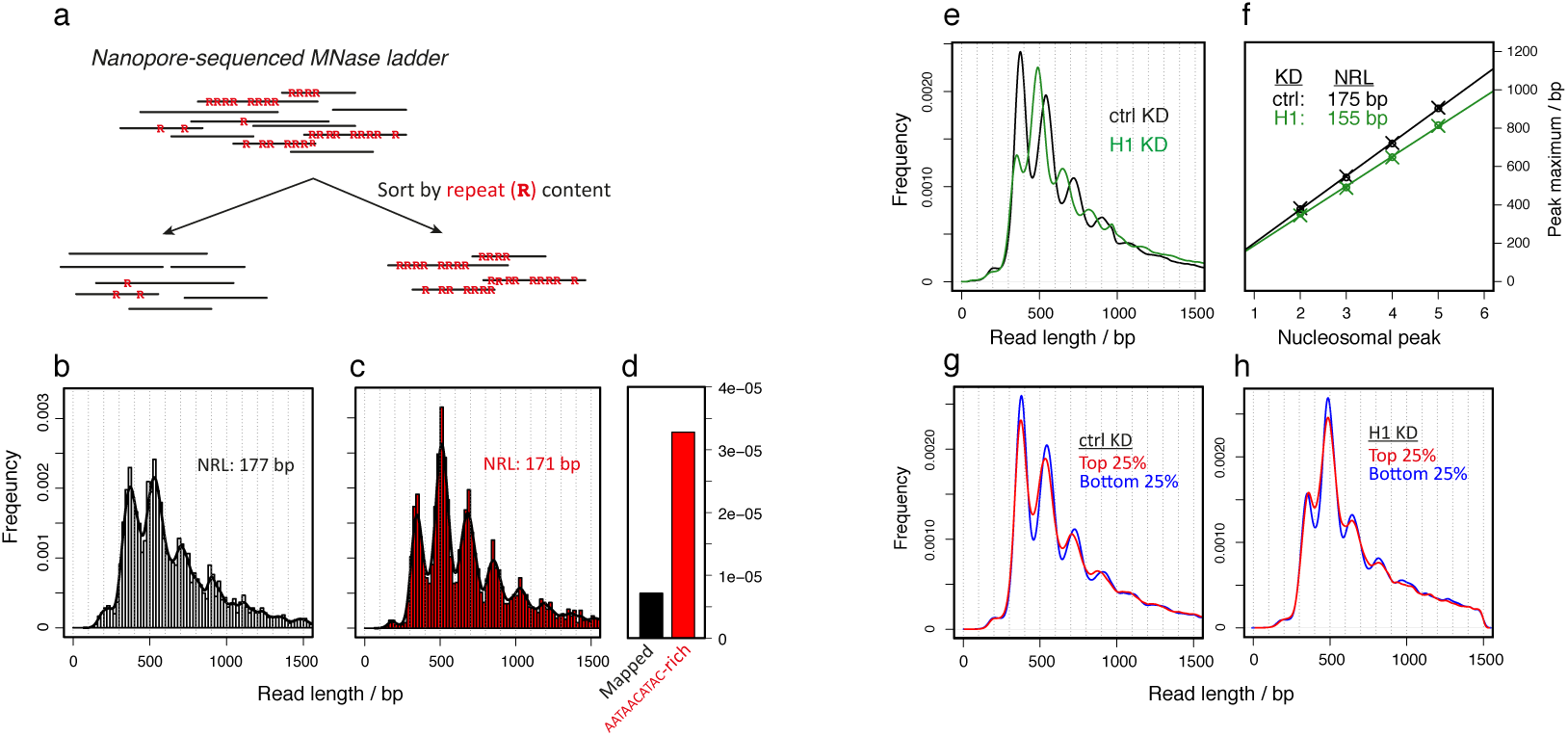
Nucleosome array regularity at heterochromatin repeats and as a function of histone H1. A, Nanopore-sequenced reads can be sorted according to the presence of sequence features (e.g. enrichment of repeat sequences), irrespective of whether they can be aligned to the mappable genome. **b**, Size distribution of 3005 randomly chosen Array-seq reads mapping to the genome (read number adjusted to fit the number of reads in **c**). **c**, Size distribution of Array-seq reads consisting to at least 30% of heterochromatic AATAACATAC repeats. **d**, Regularity scores for control and AATAACATAC-rich fragments. **e**, Size distributions of Array-seq reads derived from Kc cells depleted of H1 by RNA interference (H1 KD), or controls (ctrl). **f**, Linear regression to determine the NRL for control and H1-depleted cells. Two replicates are included for each condition, marked by circles and crosses, respectively. **g**, Size distributions for Array-seq reads mapping to either top 25% (red) or bottom 25% (blue) expressed genes in control Kc cells. **h**, Size distributions for Array-seq reads mapping to either top 25% (red) or bottom 25% (blue) expressed genes in H1-depleted Kc cells.

## H1 knockdown strongly reduces linker length

The linker histone H1 binds to the DNA entry and exit sites on the nucleosome surface and is important for the formation of higher-order chromatin structures ^31^ and a crucial determinant of nucleosome linker length ^21,32-34^. As a further test for the validity of our approach we determined how depletion of H1 in *Drosophila* Kc cells (Supp. Fig. 3a) affects the NRL. As expected, the distribution of the sequenced fragments shifted towards shorter fragment sizes in the H1 knockdown sample (Fig. 4e). This was accompanied by a considerable loss of amplitude of the dinucleosomal peak, which is probably due to the inefficient sequencing of shorter fragments in the nanopore device. Nevertheless, the peak positions could be readily used to determine the linker length in the control and H1 knockdown cells and were in good agreement with the ones obtained from an independent replicate (Fig. 4f). It shows that the NRL is reduced from 175 bp in control cells to 155 bp in the absence of H1. This difference of 20 bp fits well to earlier observations that H1 protects about 20 bp of DNA from enzymatic digestion ^35-38^. Furthermore, array regularity and linker length were reduced in highly expressed genes in Kc cells (Fig. 4g), similarly to what we had earlier observed in BG3-c2 cells. Interestingly, although the loss of regularity at highly expressed genes is still apparent in H1 knockdown cells, there seems to be no difference in linker length between expressed and silent genes (Fig. 4h). This may suggest that the reduction in linker length induced by transcription is due to a loss of H1 from chromatin, which would fit to earlier observations that active transcription reduces H1 levels in chromatin ^39-41^.

## Illumina sequencing of long fragments maps linker length with high resolution

Nanopore sequencing currently yields read numbers in the range of the low millions at best. A few thousand reads are needed to obtain a meaningful size distribution to measure regularity scores and linker length by Array-seq. Therefore, it is not yet feasible to subset the whole genome into small bins and determine array regularity and NRL for each bin. To obtain an independent measure of NRL with higher genomic resolution we explored an alternative approach. We purified the tetranucleosome-derived fragment from a gel resolving an MNase digest of BG3-c2 chromatin and subjected the purified DNA to paired-end sequencing on an Illumina HiSeq1500 sequencer (Fig. 5a). In this application, the clustering of long fragments in the flow cell is much less efficient than that of shorter DNA, therefore only samples with equivalent (long) fragments sizes can be sequenced together in the same flow cell. Indeed, the size distribution of the paired-end reads revealed that trace amounts of short fragments that contaminated the isolated tetranucleosomal DNA clustered so much more efficiently in the sequencer that they comprised a considerable fraction of all sequenced fragments (Fig. 5b). Nevertheless, sufficient numbers of tetranucleosome-size reads were obtained for the analysis of NRL. To demonstrate the usefulness of our approach we sequenced two replicates of BG3-c2 control cells and of cells depleted for the chromatin remodeler Iswi (Supp. Fig. 3b). The ATPase Iswi is part of several chromatin remodeling complexes ^42-46^ and has been implicated in the formation of regular chromatin and to influence linker length ^43,44,47^. Therefore, it seemed a good candidate to test for changes in nucleosome spacing. Indeed, whereas tetranucleosomal fragments from control cells had an average length of 663 bp, this was reduced to 629 bp in Iswi knockdown cells. We do not know how much trimming by the MNase occurs at the fragment ends, but if we consider the outer linkers are trimmed down completely, we can obtain the linker length by subtracting 4*147 bp from the total length and dividing by 3. For the control cells, this yields an average linker length of 25 bp, which is very close to the value obtained by nanopore-based Array-seq (26 bp, Fig. 1e). In Iswi knockdown cells, the linker length was reduced to 14 bp. Iswi promotes association of H1 with chromatin in *Drosophila* ^48^, so it may very well be that the shortening of the NRL we observe in Iswi-depleted cells is due to a reduced incorporation of H1. We then used the sequenced fragments to determine linker length along the mappable genome in 5000 bp bins (Fig. 5c). This revealed that different parts of the genome have different linker length. It also showed that linker length is reduced globally in cells lacking Iswi, but that different parts of the genome are affected to different degrees. By mapping the changes in linker length between Iswi knockdown and control cells, we may trace the remodeler’s activity along the genome (Fig. 5c, bottom track).

**Fig. 5.**
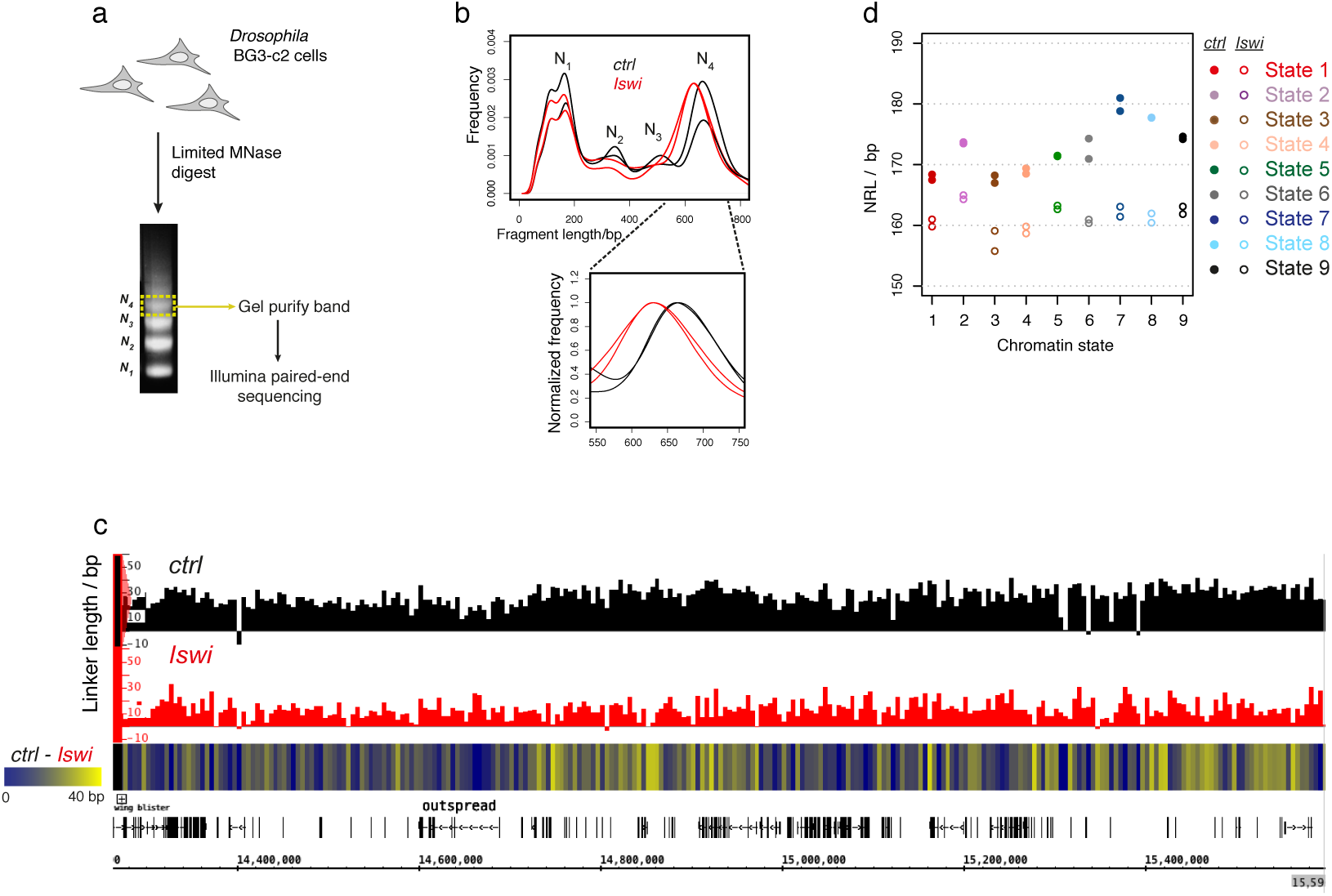
Illumina-based sequencing of long fragments allows determining the NRL along the genome. **a**, Overview of an Illumina-based Array-seq experiment. **b**, Size distributions of sequence reads derived from the purified tetranucleosomal band from BG3-c2 cells either treated with control or Iswi dsRNA (two replicates each). Note the contaminating shorter fragments. The bottom panel shows a zoom of the tetranucleosomal peaks normalized to the maxima. **c**, Genome browser screen shot showing average linker length (derived as [N_4_ fragment length– 4 × 147]/3) mapped for 5000 bp bins along the genome for control (ctrl) and Iswi knockdown cells. The heat map (bottom) shows differences in linker length between control and Iswi-depleted cells, which may indicate regional variation in the remodeler’s activity. **d**, NRL for the different chromatin states as determined by Illumina-sequencing of tetranucleosomal fragments.

Mapping the tetranucleosomal reads to the 9 chromatin states revealed considerable differences in NRL in control cells (Fig. 5d), with active promoters and enhancers (state 1 and 3, respectively) having, on average, the shortest linker length (around 20 bp), and heterochromatin and silent domains (states 7,8,9) the longest ones (around 30 bp). Interestingly, NRL length is not uniformly reduced in Iswi knockdown cells, as the states with longer linkers in normal conditions (heterochromatin and silent domains) show a particular strong decrease in NRL in the absence of Iswi. This demonstrates that paired-end sequencing of long MNase fragments is a simple yet powerful tool to map linker length with high resolution throughout the genome even in the absence of well-positioned nucleosomes.

## Discussion

We developed a new strategy to measure array regularity and NRL independently of nucleosome phasing in a genome-wide manner. We achieved this by combining classic limited MNase-digestion of chromatin yielding oligonucleosome-size DNA with nanopore-based sequencing technology. The experiment is straightforward, as the purified DNA after MNase digestion can be directly used for the simple library preparation for sequencing on a nanopore device. The lack of a PCR step during library preparation prevents the introduction of amplification biases, but also requires a minimal amount of DNA in the range of 0.1-0.5 µg per sample, amounts that can be readily obtained from cells or tissues. We used Oxford Nanopore MinION devices for all our sequencing, yielding read numbers in the 100,000s, which was sufficient for addressing our questions in the context of the *Drosophila* genome. If a larger number of reads is required, the Oxford Nanopore PromethION system may be used, which is based on the same technology but contains multiple flow-cells, each of which has about six times the sequencing capacity of one MinION device. In our case, we used Array-seq to determine patterns of chromatin regularity in different genomic ranges that were defined by chromatin states or transcriptional activity. The final goal would be to sequence deep enough to be able to subset the entire genome into small bins to determine a regularity score for each bin. To do this for the *Drosophila* genome, we estimate to need at least 10^8^ reads per experiment. However, given the rapid evolution of nanopore sequencing technology such applications may well become feasible before too long. In the meantime, paired-end Illumina sequencing of oligonucleosomal fragments allows the mapping of NRLs onto the genome with high resolution. For this approach, one has to choose whether to sequence di-, tri-, or tetranucleosomal fragments. There is a trade-off between ease of library preparation (which works best for dinucleosomes) and the sensitivity to detect changes in linker length, which is better for longer fragments as they contain multiple linkers per molecule.

A very interesting feature of the Array-seq approach is that also genomic regions that cannot be mapped to the reference genome can be analyzed. We used it to determine regularity in regions that are enriched for a classic heterochromatic satellite repeat. We found the arrays in these regions to be exceptionally regular, which is in good agreement with earlier findings of high regularity in *Drosophila* heterochromatin ^12,13^. This approach can be easily extended to any type of repeat sequence or the study of transposable elements with poorly defined insertion sites.

In general, we observed that nucleosome arrays of transcriptionally silent domains were more regular than average. It is difficult to say whether array regularity contributes to efficient silencing or is a mere consequence of it. However, we describe elsewhere that there is widespread derepression of silent genes in mutant flies lacking the regularity-promoting ISWI-type chromatin remodeler ACF ^49^. This suggests that regular nucleosomal arrays may indeed contribute to the repressive ground state of chromatin.

Maybe our most interesting finding concerns the regularity of nucleosome arrays just downstream of transcription start sites. Traditionally, the strong nucleosome phasing at the 5’ end of expressed genes observed in MNase-seq experiments is regarded as a paradigm for regular chromatin ^6,7,14,15,25,26^. Inactive genes do not show regularly phased nucleosomes at these positions. We now show that nucleosome arrays downstream of inactive transcription start sites are considerably more regular than the ones at active promoters, albeit not phased with respect to the DNA sequence. The regularity of nucleosomes with poor translational positioning is only revealed if the relation between neighboring nucleosomes is determined within single DNA molecules, as it is achieved by Array-seq. This demonstrates that the method reveals fundamental aspects of local chromatin structure that, so far, have not been accessible to genome-wide approaches. In the future, it will be interesting to test potential effects on array regularity in mutants for proteins involved in chromatin regulation and structure, particularly ATP-dependent chromatin remodelers, whose binding patterns are often difficult to determine by ChIP-sequencing ^50,51^ and whose actions could so far mostly be mapped only on predefined regions with well-positioned nucleosomes ^52-55^.

## Methods

### Cell culture and RNAi

BG3-c2 and Kc cells were cultured at 26°C in Schneider’s *Drosophila* Medium (Gibco) including 10% fetal calf serum (FCS), Penicillin-Streptomycin (and for BG3-c2 cells also 10 µg/ml human insulin). For *ISWI* knockdown, 1.5 × 10^7^ BG-c2 cells were resuspended in 4 ml medium (containing antibiotics and insulin but no FCS), seeded in a T-75 flask, and incubated with 50 µg of *in vitro* transcribed dsRNA (MEGAscript T7 Transcription Kit, Ambion). After gentle shaking for 45 min at room temperature, 16 ml of medium (+FCS/+antibiotics/+insulin) were added to the cells. Cells were harvested after 6 d. Two million cells were set aside and frozen/boiled thrice in 100 µl urea buffer to test for *ISWI* knockdown efficiency. The remaining cells were processed for MNase digestion. H1 knockdown in Kc cells was done similarly, but after 3 d the cells were harvested and the dsRNA treatment repeated on 1.5 × 10^7^ of the harvested cells. After another 3 d, the cells were harvested and processed for MNase digestion.

For knock-downs of H1 and Iswi, templates for *in vitro* transcription were amplified from *D. melanogaster* genomic DNA with the following primers:

f-H1:TAATACGACTCACTATAGGGgatgtctgattctgcagttgc
r-H1: TAATACGACTCACTATAGGGgggcttcgactttatgattcc
f-Iswi: TAATACGACTCACTATAGGGtcagagccgtctgccttattg
r-Iswi: TAATACGACTCACTATAGGGaattaaacacatcgggcagc

### MNase digestion of chromatin

1−2 × 10^8^ cells were harvested by centrifugation (500 × g, 5 min), washed in 15 ml cold PBS, and resuspended in 1 ml lysis buffer (15 mM HEPES pH 7.6, 0.1 mM EGTA, 1 mM EDTA, 0.15 mM spermine, 0.5 mM spermidine, 1x Protease Inhibitor cocktail (Roche)). After 1 min incubation on ice, 176 µl of 10% NP-40 was added, gently vortexed, and the cells again incubated on ice for 5 min. The released nuclei were spun down at 4°C and 1000 rpm (100 × g) for 10 min. They were then washed with 1 ml MNase buffer (10 mM Tris-HCl pH 7.5, 15 mM NaCl, 60 mM KCl, 2 mM CaCl_2_, 0.15 mM spermine, 0.5 mM spermidine) and centrifuged again (4°C, 100 × g, 10 min). The nuclei were resuspended in 1 ml MNase buffer and distributed into five aliquots of 200 µl. Different amounts of MNase (Sigma-Aldrich) were added to the five tubes. The exact amounts depend on MNase batches and conditions and should be determined empirically. In a typical experiment, we prepared a stock solution of 1.2 × 10^−3^ U MNase/µl and added 1, 2, 4, 8 or 16 µl of this stock solution to the 200 µl aliquots. The chromatin was then digested for 20 min at 37°C. Digestion was stopped by transferring the tubes to ice and adding 5 µl 0.5 M EGTA. To digest proteins, 12 µl of 10% SDS and 100 µg of Proteinase K (Bioline) were added and the chromatin was incubated at 50°C for 1 h. After nucleic acids were extracted with 1 volume of phenol-chloroform-isoamyl alcohol mixture (PVA, 25:24:1, Invitrogen), 50 µg RNase A was added to digest RNA at 37°C for 30 min. DNA was extracted with 1 volume of PVA. DNA was precipitated in presence of 10 µg glycogen (Roth) by adding 1/10 volume of 3 M NaOAc and 2.5 volume. EtOH and incubation for 1 h or overnight at −20°C. The precipitated DNA was pelleted at 4°C and 16,000 × g for 30 min and washed with 300 µl 70% EtOH. Pellets were shortly dried (5 min) and dissolved in 21 µl TE buffer. 1 µl of each sample was run on an Agilent Bioanalyzer 1000 DNA chip to determine degree of digestion.

### Nanopore sequencing

Partially MNase-digested chromatin retaining contiguous oligonucleosomes was used for generation of libraries for nanopore sequencing. 1 µg of purified DNA was end-repaired and A-tailed with the NEBnext UltraII end repair module (New England Biolabs, Ipswich, USA), purified with Ampure XP beads (Beckman-Coulter, Brea; USA) and ligated to AMX adapter from the Oxford Nanopore 1D sequencing Kit (SQK-LSK108, Oxford Nanopore Technologies, Oxford, UK) using NEB Blunt/TA ligase mastermix. Excess adapters were removed by adding 0.5 volumes of Ampure XP beads and washing with ABB buffer from the 1D sequencing kit. Purified library was eluted and loaded to a MinION R9.4 flowcell according to the manufacturer’s instructions and run for 48 h. Resulting fast5 raw data files were basecalled and converted to fastq format with the Albacore software (Oxford Nanopore Technologies, Oxford, UK). Reads were mapped to the *D. melanogaster* genome release 5.22 with GraphMap, a mapping algorithm specifically designed to analyze nanopore sequencing reads ^56^.

### Illumina sequencing of tetranucleosomal fragments

10 µl of DNA from partially MNase-digested chromatin was run on an 11 cm 1.2 % agarose gel in 0.5x TAE buffer at 70 V and stained with ethidium bromide. The tetranucleosomal band was cut out and purified (Gel and PCR Clean Up Kit, Macherey-Nagel). 80 ng DNA was used for library preparation with the Diagenode iDeal kit. The libraries were purified with 1x AMPure beads (Beckmann Coulter) and paired-end sequenced for 50 bp on an Illumina HiSeq1500 machine. Six tetranucleosomal samples were combined in one sequencing lane without any other samples. Fragments were mapped to the *D. melanogaster* genome release 5.75 with the bowtie mapper ^57^ with default parameters except –X = 1500 to allow for long distances between the paired reads.

### Mononucleosome mapping

Chromatin was prepared and digested as described above, but the nuclei were resuspended in 600 µl of MNase buffer instead of 1 ml, distributed into three 200 µl aliquots and digested with 2, 5 or 10 µl of the MNase stock solution. Ten microliter of each digested sample were run on an 11 cm 1.2 % agarose gel in 0.5x TAE buffer at 70 V and stained with ethidium bromide. The sample containing predominantly mononucleosomal fragments (digested with 10 µl MNase stock solution) was chosen, the mononucleosomal band cut out, and the DNA purified (Gel and PCR Clean Up Kit, Macherey-Nagel). 10 ng of the DNA was used to prepare a sequencing library as described elsewhere ^55^. The library was paired-end sequenced for 50 bp on an Illumina HiSeq1500 machine. Fragments were mapped to the *D. melanogaster* genome release 5.75 with the bowtie mapper ^57^ with default parameters.

### Data analysis

Mapped reads derived from nanopore- and Illumina-sequenced long fragments were converted to .bed files with *BEDTools* ^58^ and imported into an R environment (RStudio), where statistical analysis on the size distributions was performed. To investigate the size distribution of fragments enriched for heterochromatic AATAACATAG repeats, unmapped nanopore reads where converted to .fasta format and searched for occurrences of the AATAACATAG motif with the FIMO tool ^59^ with default settings except --max-stored-scores 10,000,000. The FIMO output file was then imported into R, where only the reads with perfect motif matches were selected and kept. For each read, the number of perfect matches for the repeat motif was multiplied with 10 (the length of the motif) and divided by the read length. This yielded the motif enrichment for each read. Every read that had a motif enrichment of at least 0.3 (30% or more of the read consist of the motif) was kept and used for size distribution analysis.

For mononucleosome mapping, the mapped paired-end sequenced reads were converted to dyad coverage vectors by size-selecting fragments of length >=120 and <=200 bp and resizing their length to 50 bp fixed at the fragment center, and aligned at transcriptional start sites (*D. melanogaster* dm3 genome version).

Genes were sorted according to their expression levels in BG3-c2 and Kc cells based on data published in ^60^.

### Data deposition

Raw nanopore and Illunina .fastq files and mapped .bed files are deposited on the Gene Expression Omnibus (GEO) repository with accession number GSE110807 (www.ncbi.nlm.nih.gov/geo/query/acc.cgi?acc=GSE110807).

## Acknowledgements

This project was supported by grant Be1140/8-1 of the German Research Council (DFG) to PBB. We thank Tobias Straub, Tamas Schauer and Dhawal Jain for inputs for data processing, Angelika Zabel for help with chromatin preparations and MNase digests, and Philipp Korber for critical reading of the manuscript and many helpful suggestions to improve it.

## Contributions

S.B. devised the project, performed the experiments and analyzed the data under the supervision of and with input from P.B.B.. S.K. performed all sequencing under the supervision of H.B.. S.B. and P.B.B. wrote the manuscript.

